# Control without controllers: Towards a distributed neuroscience of executive control

**DOI:** 10.1101/077685

**Authors:** Benjamin R. Eisenreich, Rei Akaishi, Benjamin Y. Hayden

## Abstract

Executive control refers to the regulation of cognition and behavior by mental processes and is a hallmark of higher cognition. Most approaches to understanding its mechanisms begin with the assumption that our brains have anatomically segregated and functionally specialized control modules. The modular approach is intuitive: control is conceptually distinct from basic mental processing, so an organization that reifies that distinction makes sense. An alternative approach sees executive control as self-organizing principles of a distributed organization. In distributed systems, control and controlled processes are co-localized within large numbers of dispersed computational agents. Control then is often an emergent consequence of simple rules governing the interaction between agents. Because these systems are unfamiliar and unintuitive, here we review several well-understood examples of distributed control systems, group living insects and social animals, and emphasize their parallels with neural systems. We then re-examine the cognitive neuroscience literature on executive control for evidence that its neural control systems may be distributed.

## I. Introduction

Executive control refers to the brain’s ability to regulate its own processing. It coordinates multiple competing demands, controls attention, gates working memory, and encodes and retrieves long-term memories. It also maintains and switches task set, inhibits disadvantageous actions, and regulates the explore/exploit tradeoff and curiosity (Miller & Cohen, 2001; Shiffrin & Schneider, 1977; Braver & Barch, 2006; Cole & Schneider, 2007; Miller, 2000a; Ridderinkhof, van den Wildenberg, Segalowitz, & Carter, 2004; Kidd & Hayden, 2015). Understanding executive control is critical for understanding self-control and its failures (Aron, Robbins, & Poldrack, 2014; Knoch & Fehr, 2007; Hare & Rangel, 2009). More broadly, failures of executive control are hallmarks of many diseases, including addiction, depression, and obsessive-compulsive disorder, and successful treatments of these diseases often target executive control (e.g. Milad & Rauch, 2012; Ursu et al., 2003; Volkow & Fowler, 2000; Kalivas & Volkow, 2005).

A brain can be understood as a *control system*, a collection of interacting components within an organizational structure that produces adaptive actions based on information about the current state of the internal and external worlds (Pezzulo & Cisek, 2016; Gallistel, 2013; Lashley, 1951). As we process sensory inputs and generate actions, the brain monitors that processing and, if it detects the need to change, it regulates it. But how is executive control in the brain implemented by the interactions of its constituent parts, individual neurons?

### Modular and distributed control systems

The standard approach to understanding control starts with the assumption of modularity. In a modular control system, regulation is derived from a central controller, which is a discrete subsystem with a specialized function. In a modular system, it is theoretically possible to draw a line through anatomical space separating localized control regions or circuits (often the prefrontal cortex and striatum) from more basic processing (caudal cortical) regions (Botvinick et al, 2001; Miller & Cohen 2001; Miller, 2000). This specialization means that control regions (or networks) regulate, but do not participate in, the underlying stimulus-to-action transformation processes (Figure 1). Such a view is consistent with a long tradition emphasizing the brain’s modular architecture (Fodor, 1983; Minsky, 1988; Kanwisher, Mcdermott, & Chun 1997; Bertolero, Yeo, & Desposito 2015). But it is not the only possible view.

**Figure 1.**
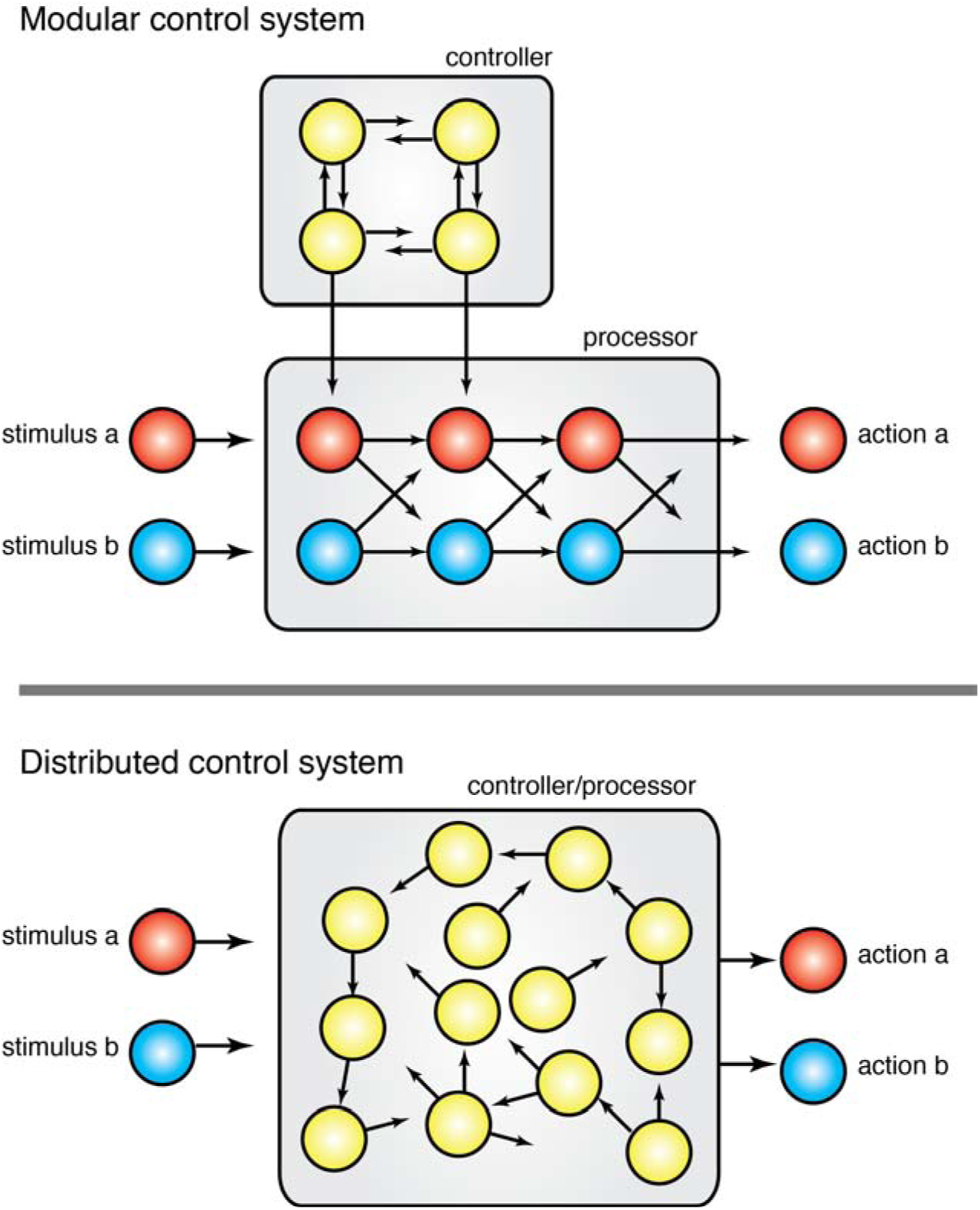
Contrasting organizations of modular and distributed control systems. Within modular control systems, processing and control elements are distinct and localized to specific areas. By contrast distributed systems combine control and processing elements, often into individual agents.

The alternative approach envisions executive control as distributed processes in which there is no dedicated and specialized controller (Figure 1). Instead, in a distributed control system, regulatory functions are dispersed across a large number of individual elements or carried out by the interaction among them (Couzin, 2009; Sumpter, 2006). In most such systems control elements are co-localized with processing elements, and those elements have somewhat autonomous function. For this reason they are often called *agents*. These agents (or any other individual elements in a distributed control system) sense the properties of their local environment and adjust their own behavior based on simple rules. Agents normally have no knowledge of the overall state of the system, and the response of the system as a whole is often qualitatively dissimilar from those of the elements. In other words, in such systems, control is often an *emergent* function (McClelland et al., 2010; Hofstadter, 1985, Ch. 25; Mitchell, 2009).

The distributed viewpoint derives inspiration from early studies on cybernetic, connectionist, and parallel distributed processing models (Rummelhart et al., 1988; Weiner et al., 1944; Grossberg, 1974; Hopfield, 1982). As noted in a review of the topic by Botvinick and Cohen (2014), the connectionist heyday of the late 70’s and early 80’s coincided with the development of formal ideas of control (Posner & Snyder, 1975; Shiffrin & Schneider, 1977; Norman & Shalice, 1986; Baddeley & Hitch, 1974). It is ironic then that almost all models of executive control, even relevant PDP models, take as given that control is functionally and anatomically modular (Botvinick & Cohen, 2014). Nonetheless, history has vindicated this approach: the modular idea is well supported by empirical data. Specifically, neuroscientific research consistently points to dorsal prefrontal structures (especially the dorsal anterior cingulate cortex, dACC, see below), as well as superior parietal cortex and parts of the brainstem as the brain’s control system (Holroyd & Coles, 2002; Botvinick & Cohen, 2014; Ridderinkoff et al., 2004; Shenhav, Botvinick & Cohen, 2013; Miller & Cohen, 2001; Sleezer & Hayden, 2016; Floresco, 2015; Mansouri et al., 2007).

### Revisiting the distributed processing view

Still, we believe that it is time to revisit a distributed approach to control. Several factors motivate this belief. First, our understanding of the neuronal (i.e. single unit) responses of the putative executive regions is only now maturating. Some of this work emphasizes the broad overlap in functions of the prefrontal and posterior regions; these functions appear to include both processing and executive roles (Cisek & Kalaska 2010; Kim & Shadlen, 1999; Chafee & Goldman-Rakic, 1998; Postle, 2006; Awh & Jonides, 2001; Sleezer & Hayden, 2016a; Sleezer, Castagno, & Hayden, 2016). Second, new anatomical and functional techniques emphasize the fundamentally non-modular organization of the brain (Misic & Sporns, 2016; Wang et al., 2015; Farah, 1994; Kristan & Shaw, 1997; Plaut, 1995). Third, major recent advances in computation have come from abandoning classic (GOFAI)-style symbol manipulating systems in favor of deep learning algorithms that are distributed and recurrent (e.g. Lecun, Bengio, & Hinton, 2015; Hinton & Salakhutdinov, 2006). These approaches highlight the power and flexibility of non-modular network organizations. Finally, recent years have seen a greater understanding of the mechanisms of distributed control in non-brain biological systems, leading to a greater appreciation of the strengths and of the biological plausibility of such systems (Couzin, 2009; Passino, Seeley, & Vischer; 2007).

Reified models of executive control – in which conceptual elements like monitor, controller, and processor have direct correspondence with neuroanatomy – are intuitive. But distributed models are less so. To mitigate this problem here we offer a summary of the basic principles of distributed control systems, with an emphasis on natural examples.

## II. Principles of distributed control systems

### Principle 1: Horizontal information flow

Within a modular control system, information flows linearly from lower level processing units to the controller. By contrast information flow within distributed systems is characterized by horizontal communication between adjacent members. In other words, information is derived from neighbors, not from a central communicator. Consequently, no single member of a distributed system is knowledgeable about the entire system. Each member can know what their neighbor is doing, and possibly what their neighbor knows, through localized interactions.

A good example of information flow within a distributed system is a herd of baboons on the move (*Papio anubis,* Couzin &Krause, 2003; Strandberg et al, 2015). Even though they have a hierarchical dominance system, no single member of the troop knows for sure where to go but several members have some limited and likely noisy knowledge (Figure 2). The wisdom of the crowd is better than any individual’s guess, as in many collectively moving animals – including humans (Codling, Pitchford, & Simpson, 2007; Simons, 2004; Hamilton, 1967; Bergman & Donner, 1964; Walraff, 1978; Mallon, Pratt, & Franks, 2001; Conradt & Roper 2003). The baboon troop thus uses a collective decision-making strategy. Individuals begin to head off towards their best guess and as they do this, troop members compute the average of the members they observe. Unlike in a modular system each member may be simultaneously a decision maker and a data point for other decision makers.

**Figure 2.**
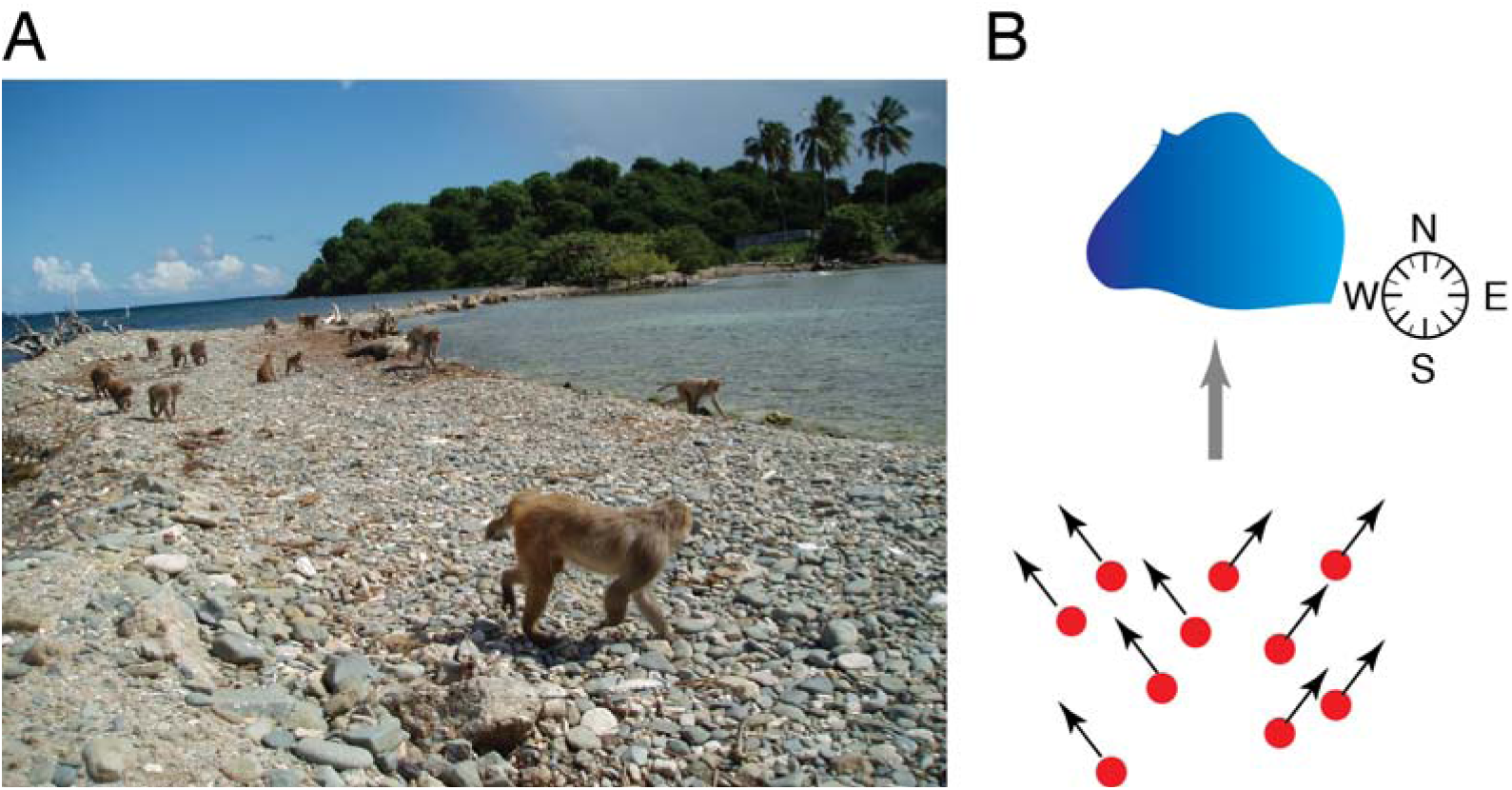
Group movement strategies often illustrate the principle of horizontal information transfer. **A.** Rhesus monkey troops on Cayo Santiago migrate multiple times each day and may use distributed consensus procedures to choose a direction. **B.** Cartoon birds eye view illustrating split voting situation. If the troop is split between a northeast and a northwest direction, the consensus will not be the average (north) but one of the two modal directions.

Normally this strategy leads efficiently to a rapid consensus (Couzin & Krause, 2003; Conradt & Roper, 2003). In cases where there are two different modal preferences – say, when northeast and northwest are both good directions but true north is not, this averaging strategy leads to a suboptimal choice (Figure 2B). For this reason individuals should be – and are - sensitive to bimodal distributions among the group and, in that case, randomly choose one of the two modal directions (Strandberg et al, 2015). Similar patterns are observed in pigeons and human crowds (Biro et al., 2006; Dyer et al, 2008).

In this example, the input is the environmental clues (including memories) about the best direction to head and the output is a group path. Information is distributed across individual troop members who communicate locally with each other. Drawing from the local interactions among members, the group chooses a better output than all the constituent individuals. The decision is also controlled in a closed-loop manner: the group can monitor its own performance (it can detect split voting) and regulate its voting strategy (averaging to bifurcation-then-averaging), even though no individual serves as the specialized monitor or regulator. Instead, monitoring and control proceed through local, horizontal connections between members.

The idea of horizontal flow of information from adjacent members is also often a description of neuroanatomical organization. Neurons, like troop members, tend to have limited view of the activity of the whole, limited ability to communicate with the whole, incomplete information, no knowledge of the larger factors that determine the group’s well-being, and no obvious leadership. However, neurons do have a rich network of connections to adjacent neighbors and cortical areas that supports a localized flow of information. While the brain also has centralized global signaling, in the form of neuromodulators (and possibly cortical oscillations), the bandwidth of these signals is limited and the timing may be too slow to affect on-line decision processes. Similar to a baboon troop, the information gained from equal and adjacent members has a large effect on the regulation of its neural function.

### Principle 2: Stigmergy

In the case of the baboons, it is notable that the control signal is the movement of neighbors. Thus, in a strongly non-modular way, the control signal is precisely the output of the underlying process (also movement of individuals). It is a *stigmergic* system (Bonabeau, Dorigo, & Theraulaz, 1999; Theraulaz, Bonabeau, & Deneubourg, 1998; Couzin, 2009).

A familiar example of stigmergic signaling is lawn shortcut generation on college campuses. A student following the trod path also – weakly but surely – strengthens it (Figure 3.). Another example is pheromonal trails in foraging ants (Hölldobbler & Wilson, 1990; Wilson, 1971). As a scout forages she lays a scent that other scouts will follow to valuable food sources. The scent evaporates quickly, so rich food patches, which attract many ants, will have stronger paths leading to them. An ant that, by chance, discovers a shortcut will produce a trail with a stronger scent (because, being shorter, it takes less time to traverse and thus has more scent, Beckers & Deneubourg, 1992). In this way, pheromones allow ant colonies to find rich food sources and develop shortest path routes without any centralized control (Aron, Beckers, & Deneubourg, 1993; Jackson & Chaline, 2007; Beekman, Sumpter, & Ratnieks, 2001).

**Figure 3.**
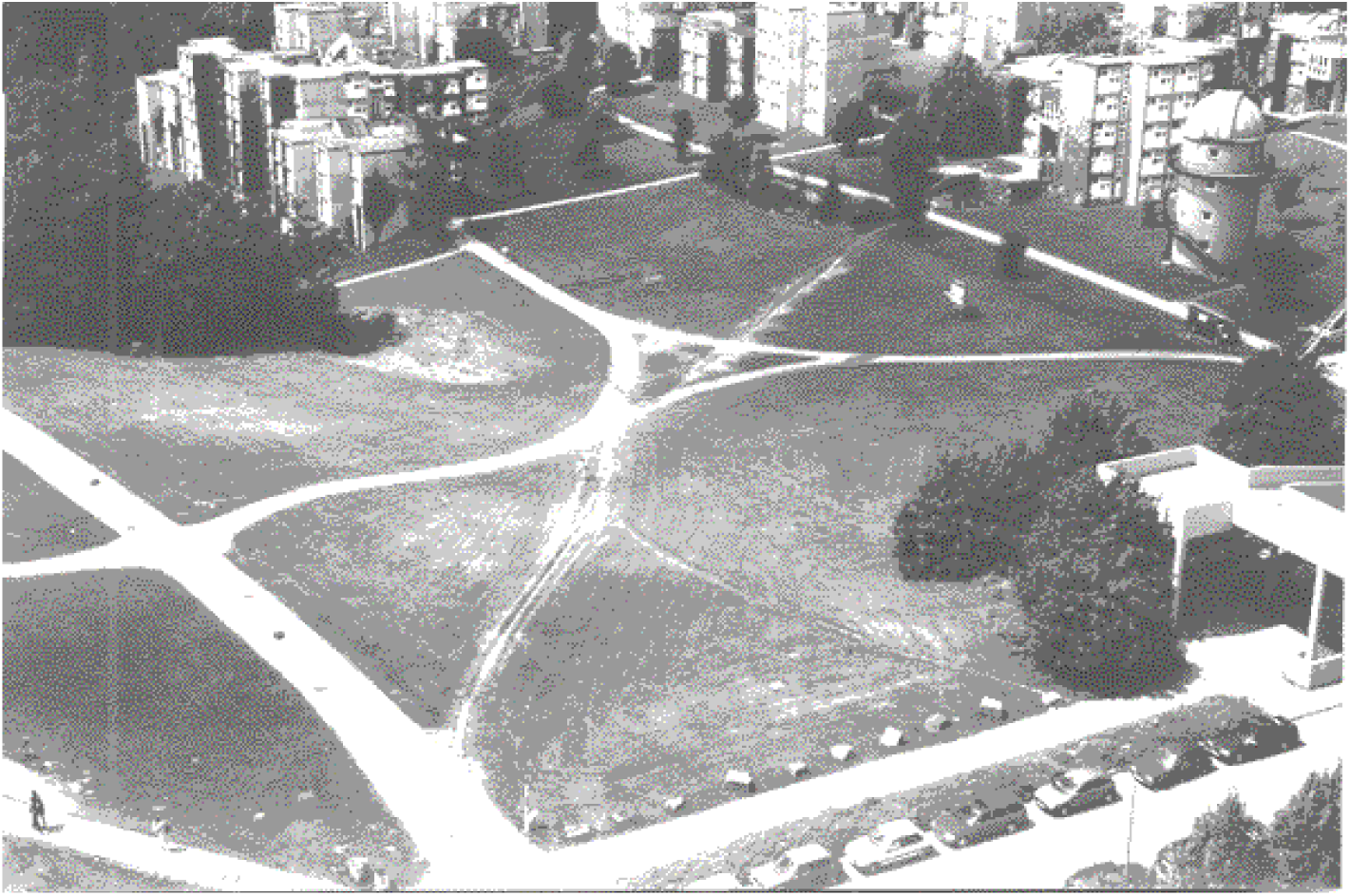
Humans can collectively identify, create, and maintain efficient paths across lawns on college campuses. Reproduced with permission from “Modeling the evolution of human trail systems” (Helbing, Keltsch, & Molnar, 1997).

Another example of stigmergic control comes from the process of neural differentiation of sensory organ precursors within the developing fly brain (*Drosphilia melanogaster*) (Navalakha & Bar-Joseph, 2011). During development some cells within the neural clusters of the fly brain become sensory organ precursors (SOPs); these cells form the backbone of the sensory system later in development. Determination of which cells become SOPs follows an algorithmic process that produces a maximally independent set distributed throughout the brain. Functionally each cell will propose itself as a possible SOP. If any neighboring cell has already become a SOP the proposing cell will not differentiate. As a consequence of this process the likelihood of an unconnected cell differentiating increases with time (Afek et al., 2011; Navalakha & Bar-Joseph, 2011). By using information about the structure of neighboring cells, each cell is able to differentiate appropriately so that the whole brain achieves an equal spacing of sensory organ precursors. The brain cells do this rapidly and without the need for a monitor or knowledgeable controller sending distinct control signals. All the monitoring and control that is needed occurs locally, within each cell.

Principles of stigmergy within executive control processes relates to neural function quite directly. Neurons produce chemical outputs that modulate responses of downstream neurons. These outputs are both the computational outputs of the neurons and a way to modulate activity of their neighbors. In the short term, excitatory and inhibitory outputs increase and reduce, respectively, the likelihood that the target will fire. In the long term, activity (especially coincident activity) promotes synaptic plasticity thus up- or down-regulating that target’s firing on longer timescales. Within cortical regions, these localized interactions could very well lead to emergent control signals without the need for a dedicated controller (Couzin, 2009).

### Principle 3: Feedback loops

Feedback is a powerful tool in any dynamical system. It can have positive effects. When fish school, a few peripheral individuals may detect a potential predator and turn away from it (Treherne & Foster, 1981; Couzin & Krause, 2003). Neighbors who follow an average-direction rule then turn and also affect their neighbors, the effect multiplies, and the traveling wave of turning fish turns the whole school away. The amplification protects many more fish than were able to detect the predator. Similarly, feedback loops are a mainstay of other distributed leaderless systems; even audience clapping, for example, can depend on feedback effects (Néda et al., 2000)

However, feedback loops can be dangerous as well (Giraldeau & Valone, 2002). Simple effects can snowball and, because the system is distributed, there is no central controller to stop it. For example, ants leaving a pheromonal trace can find their own trail, and start going in a circle – a literal feedback loop called an ant mill (Delsuc, 2003). Another important example of a feedback loop is a marketplace bubble (Porter & Smith, 1994; Smith, Suchanek, & Williams, 1988). If a speculator believes a commodity will go up in price, she may bid a slightly greater price than the current one. This bidding will serve as a signal to other investors that the commodity may be a wise investment. As they bid up the price, their initial assessment will be proven to be right, and other investors will gain interest. This pattern can lead to runaway prices, but only up to a point; as soon as this point is reached, the price will crash.

The tendency to boom and bust can lead to market instability and to underinvestment. In marketplaces, centralized control (such as trading limits) can solve these problems. Without that kind of control, avoiding these kinds of malign feedback loops requires careful calibration of the rules each individual follows. Such calibrations often involve complementary negative feedback loops (Grünbaum, 1998). The analogy to brains, which have many overlapping positive and negative feedback loops, is quite direct.

### Principle 4: Self-organization through simple rules

Many distributed control systems are self-organized (Sumpter, 2006). Classic examples of self-organization include bird flocks and fish schools (Aoki, 1982; Couzin, 2009; Reynolds, 1987). No leader bird rallies its mates and tells them where to fly; nor does a leader monitor the flock and guide its performance like coxswain on a crew team. Instead, the structure of the bird flock is a consequence of several simple principles followed by all individuals. These include rules about distance between adjacent birds (not too far and not too close, more or less) and rules about when to turn (follow the group average, Couzin & Krause, 2003). The specific rules, not a leader-bird, determine the shape of the flock (Figure 4).

**Figure 4.**
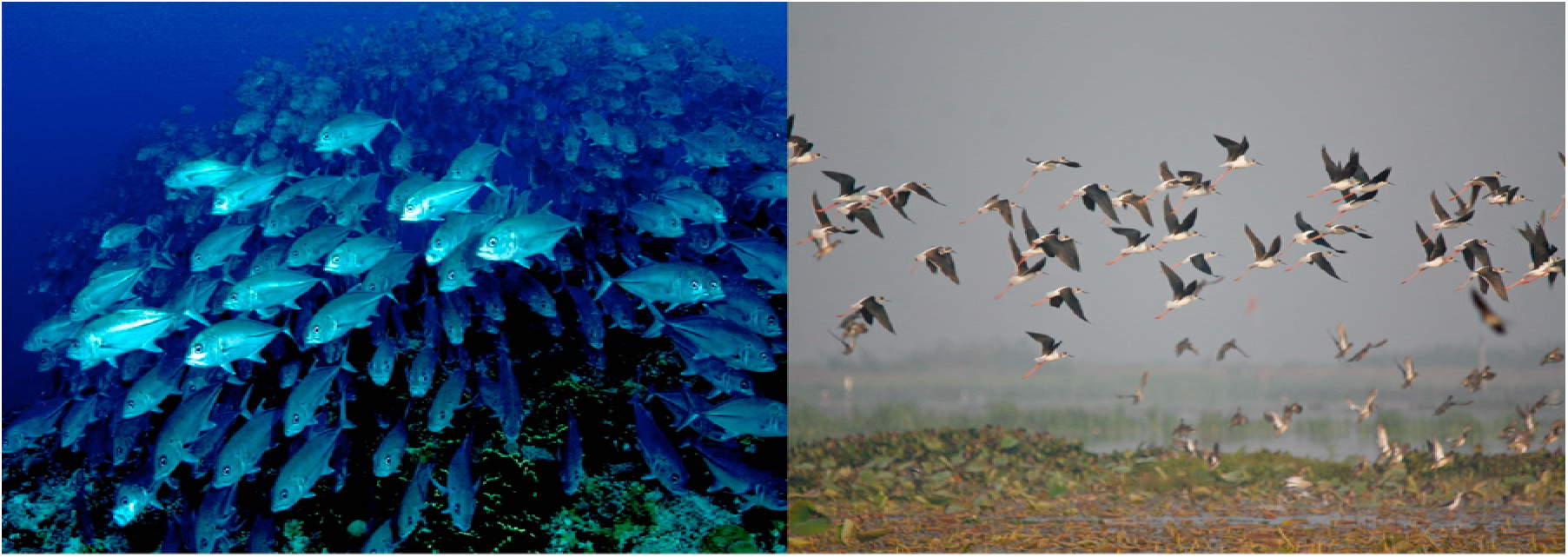
Simple rules of distance and spacing determine the shapes of both fish schools and bird flocks. [Fish picture: Gordon Firestein - Seacology USA, Bird Flock: Faisal Akram]

Self-organization is an appealing principle because it is easy to implement and is robust to degradation (Sumpter, 2006). In contrast, the centralized systems are vulnerable to the loss of the controller: Remove a switch and the whole railyard breaks down; remove the coxswain and the rowers start hitting each other’s oars; remove one bird and the flock swiftly adjusts. Self-organization also allows complex adaptive behavior without programming expensive control systems. Self-organized systems can be “fast, cheap, and out of control” (Brooks & Flynn, 1989). These features – ease of implementation, graceful degradation, and robustness, makes it appealing for analyzing neural systems. One well-known example of a self-organizing system in neurons is central pattern generators, in which the activity of the ensemble is an emergent product of the interactions of the elements, none of which follows the pattern in miniature.

This does not mean all distributed systems are leaderless. There are many contexts in which formation of leadership is favored (Couzin et al., 2005; Fischoff et al., 2007; Dyer, 2009; Robson & Traniello, 1999; Reebs, 2000). Dominance hierarchies and other leadership structures are selected in many species, although leadership is seldom absolute. And there are intermediate cases - even in the case of baboons, some individuals are recognized as having greater knowledge of the right path and their opinion is more highly weighted (Strandburg et al., 2015).

Presumably, we can classify control systems on a spectrum from fully distributed and leaderless to strictly segregated and hierarchical; the specific organization observed for any system will depend on the environment in which it evolved. This fact is important to remember when considering neural systems, which may have some specialization of function (Botvinick et al., 2001; Rougier et al., 2005; Kanwisher, Mcdermott, & Chun, 1997).

### Principle 5: Quorum-sensing

Agents in distributed systems have very limited field of view in their monitoring capabilities. In other words, it is often difficult to see the forest for the trees. But sometimes it is critical to see the forest to make the best decision. In these cases, agents must engage in *quorum-sensing*: a type of consensus-based control mechanism wherein a set threshold or quorum determines the course of action (Mitchell, 2009).

There are many mechanisms for quorum-sensing; what unites them is that they do not require centralized control. For example, bacteria can produce diffusible chemicals (which can serve as a type information) and chemical concentration in the environment gives a measure of quorum (Waters and Bassler, 2005). One critical feature of any consensus-seeking measure is that it must terminate; it should also do so relatively quickly. Failures to do so can be costly, as in the case of Buridan’s ass (Lindauer, 1957; Pais et al., 2013).

Often, individuals can sense the state of conspecifics in their local environment and extrapolate to an estimate of group state. Simply averaging the states of neighbors can be helpful in some circumstances, as in bird flocks and some fish schools. One study showed that an individual schooling three-spine stickleback fish (*Gasterosteus aculeatus*) can adopt a non-linear monitoring function that produces better group behavior emergently (Ward et al., 2008). Specifically, groups of fish tended to ignore information from single neighbor but responded when two fish conveyed the same information. This non-linear criterion can reduce the probability of amplifying noise but can still effectively detect signals.

The need for agents to sense the properties of the whole, or of large subgroups, is a major problem in brain systems as a whole. This problem is acute in executive control systems, which often rely on changing processing as a function of global conditions. Without holistic integrating neurons, it is difficult to imagine a direct solution to the problem. For this reason, studies of quorum-sensing systems, which solve the problem indirectly, are particularly likely to be helpful in understanding the neural basis of control.

## III. Distributed solutions to classic executive control problems

Studies of executive control tend to focus on processes for solving a familiar set of cognitive problems. Prominent among these processes are regulation of stop/go behavior, speed/accuracy tradeoffs and conflict detection and resolution (Bogacz et al., 2009; Aron, Robbins, & Poldrack, 2014; Botvinick et al., 1999; Miller & Cohen, 2001). These operations have analogues outside of neuroscience, including in distributed control systems of natural and artificial mechanisms. In this section, we investigate how some examples of distributed control systems handle these executive control problems through the fundamental elements outlined above. Other important executive functions, which we do not consider, include working memory, attention, task set maintenance and switching, regulating the balance of explore vs. exploit behavior, and aspects of reinforcement learning. Several of these have likely correlates in distributed control systems as well. See, for example, (Couzin et al., 2002; Couzin, 2009; and Passino, Seeley, & Visscher, 2007) for speculation about how distributed processing systems can implement working memory, attention, and regulation of long-term memories.

### Stopping and going: Vibrio fischeri bacteria

Initiation and inhibition of behavior is a simple and important executive function (Jin & Costa, 2010; Schall, 2001; Aron, Robbins, & Poldrack, 2004; Niv et al., 2007; Hampshire & Sharp, 2015; Kacelnik et al., 2011). Coordination of these two antagonistic processes can produce both simple responses and complex behaviors. Stop/go behavior involves elements like precise timing, inhibition of prepotent responses, and control of vigor. Another important but less well-appreciated requirement is avoiding intermediate responses, so that the system can either fully stop or fully go, without drifting between the two extremes. In other words, being indeterminate can be costly and even lethal in urgent situations so that the distributed system has to be able to deal with this problem.

Our example of stop/go control in a distributed control system comes from the luminous bacterium *Vibrio fischeri* (Waters & Bassler, 2005; Nealson & Hastings, 1979; Miller & Bassler, 2001). This single-celled organism lives in the light organ of the Hawaiian bobtail squid (*Euprymna scolopes*) and emits light when the squid hunts at night. The light serves to camouflage the squid that otherwise would be visible in the form of a moonlit silhouette to prey below it (Visick et al., 2000). During the day the squid hides from potential predators in the dirt and turns its eyes off by extruding most of the bacteria into the surrounding ocean. As the day progresses the remaining bacteria reproduce rapidly, and, by nightfall, have replenished their stock so that there are enough bacteria to serve as an effective camouflage.

The control problem comes from the fact that the bacteria must not luminesce during the day as they are reproducing. Instead they need to switch to lighting at night all at once. In other words, bioluminescence needs to be both inducible and repressible (Nealson & Hastings, 1979). Because of their reproduction pattern, they can do this by waiting until there is a quorum of other *V. fischeri* bacteria in the squid light organ. But how do they know how many others there are? Quorum sensing. *V. fischeri* release a chemical known as acyl-homoserine lactone (AHL). They then measure the concentration of this chemical in their local environment by the transcription activator protein LuxR, which creates a complex that induces transcription of genes needed for luminescence (Kaplan & Greenberg, 1985; Stevens & Dolan, 1994). The transcription process is only triggered when the local density of AHL reaches a predetermined threshold, which serves as a go signal for the bacteria (Figure 5).

**Figure 5.**
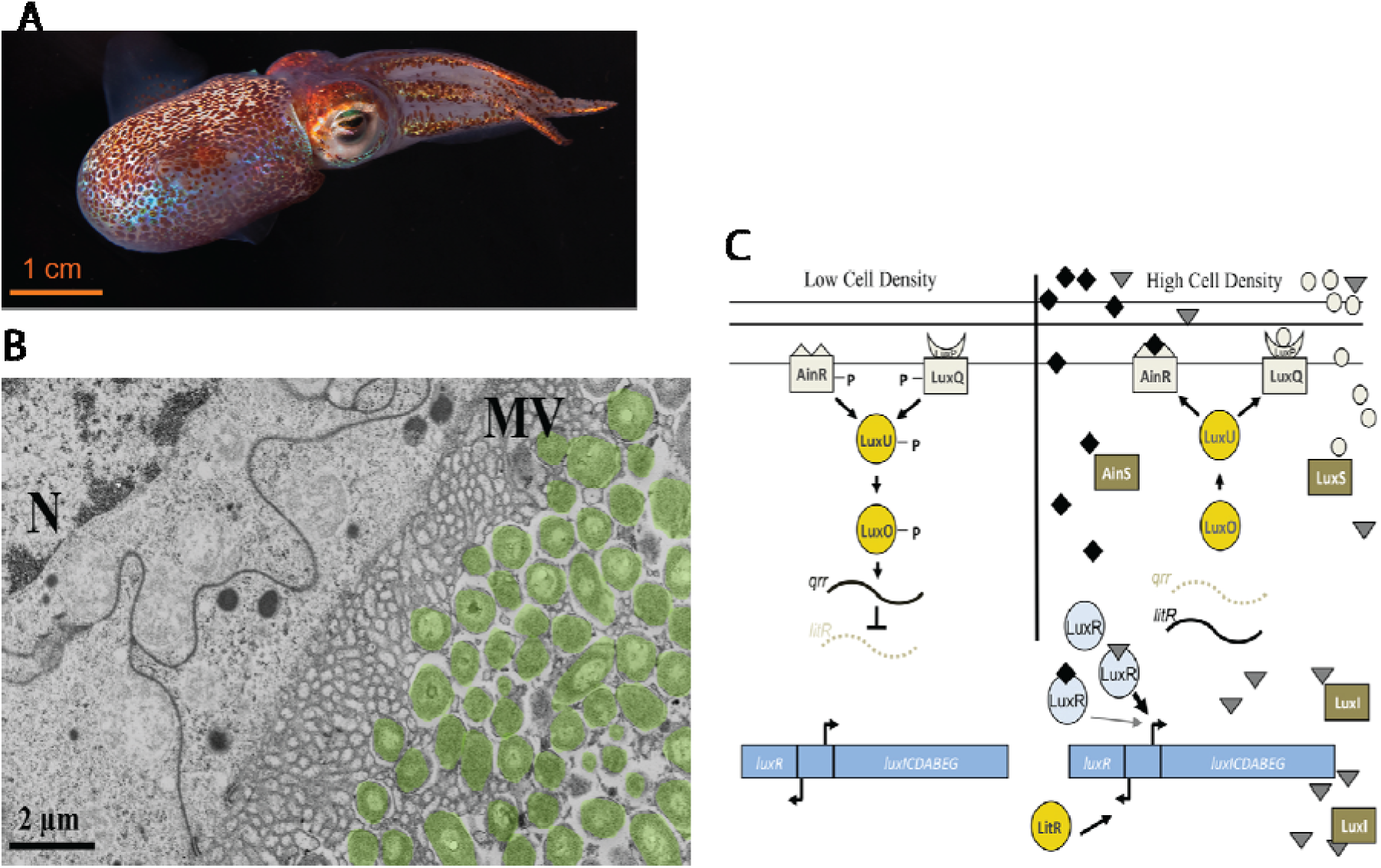
**A.** Hawaiian squid (*Euprymna scolopes*). **B.** Image of *V. fischeri* embedding into microvilli of host epithelial cells. **C.** Illustration of control circuit for regulation of luminescence through chemical detection in *V. fischeri*. Credits: (A,B) reproduced with permission from “Divining the essence of symbiosis: Insights from the squid-vibrio model.” (McFall-Ngai, 2014). (C) reproduced with permission from “Gimme shelter: how vibrio fischeri successfully navigates an animals multiple environments” (Norsworthy & Visick, 2013).

There are several features used by the system to stop, i.e. to prevent premature luminescence. These features work by implementing negative feedback (Waters & Bassler, 2005). One feature is regulation of the stability of the constituent proteins: they are more stable when AHL is more concentrated (Zhu & Winans, 1999). Another is active pumping of AHL out of the cell: this process reduces cytoplasmic levels of AHL and thus dampens sensitivity until AHL concentration is high enough to overwhelm the pumping mechanism (Pearson et al., 1999).

Several features of this stop/go process are notable here for the curious neuroscientist. First, the system implements a clock-like function by taking advantage of the consistency in reproduction rates of its own members. No member or subgroup serves as a clock or other timer function. In other words, the timing function is an emergent property of the system. Second, there is no centralized site that tells the bacteria when to glow; each individual agent makes up its own tiny mind, but, because they are in the same environment, their activity is effectively coordinated through the localized cross-signaling of individual cells. Third, the system implements a specific and precise threshold-crossing process (a simple rule based on concentration levels of AHL), even though no abstract decision variable is calculated or represented. Finally, there is no need for any kind of modular self-control or inhibition. The lack of glowing (repressability) is simply a consequence of the fact that there are insufficient concentrations of chemicals to drive the glowing; inhibition in this system is an emergent process (cf. Hampshire & Sharp, 2015).

### Speed-accuracy tradeoffs: ants

A decision made without taking the time to gather all the evidence may not be as accurate as a deliberate one, but it will have the virtue of speed (Houston, Kacelnik, & McNamara, 1982). If time is costly (as when faced by an attacking predator) it may be worth going for the first good response, but if the decision-maker has all the time in the world, it’s probably worth doing some pondering. Speed-accuracy tradeoffs are a staple of cognitive psychology (Busemeyer & Townsend, 1993; Wickelgren, 1977; Roitman & Shadlen, 2002; Chittka et al., 2003; Gigerenzer & Goldstein, 1996; Bogacz et al., 2010) and animal psychology (Chittka, Skorupski, & Raine, 2009). Like humans and animals, many distributed decision-making systems make speed-accuracy tradeoffs, including slime molds (*Physarum polycephalum*) and honeybees (*Apis mellifera*, Dussutour, Latty, & Beekman, 2010; Passino, Seeley, & Visscher 2007).

When looking for a new nest, individual ants (*Leptothorax albipennis*) leave the nest and evaluate potential locations within a few square meters (Franks et al., 2002; Franks et al., 2003). These ants prefer to live in small colonies in thin cracks in rocks and are therefore easy to study in laboratory conditions (Franks et al., 2002). An ant that finds a potential nest site will recruit other ants to evaluate it by leading a tandem run back to the site. Thus, each site is evaluated by a large number of individuals, each of whom presumably makes a worse (less accurate) decision than the cumulative choice of several ants. Unlike bees (see below) individual ants appear to evaluate and compare multiple sites, giving them more individual knowledge and requiring smaller quorum sizes (Franks et al., 2002; Pratt et al., 2002; Franks et al., 2003). If enough ants appear at a single site, scouts recognize a quorum, and the quorum catalyzes a change in their behavior; scouts now carry their nestmates to the new site and deposit them there (Pratt et al., 2002; Franks et al., 2002).

This whole search and quorum-sense process is slow but accurate. But if the situation calls for a fast decision (such as during windy weather or threat of predation), the ant colony can make a speed-accuracy tradeoff (Franks et al., 2003). Specifically, each ant can reduce the threshold it uses to decide whether to switch from tandem run recruitment mode to carrying mode. The tandem run, being slower, allows other ants more time to discover other sites; the carry terminates the process more quickly. The ant itself doesn’t know explicitly about the speed-accuracy tradeoff; it just has an internal sense of weather and adjusts its quorum-sensing procedure – and the group’s speed-accuracy tradeoff is an emergent consequence (Franks et al., 2003).

The neuroscience of the speed-accuracy tradeoff is not fully understood, but the parallels are easy to discern. It is believed that there is a threshold integration process for perceptual decisions (Bogacz et al., 2010). Recent work suggests it may involve changes in the baseline activity of neurons that serve as cortical integrators that bring them closer to threshold (Ivanoff, Branning, & Marois, 2008; Van Veen, Krug, & Carter, 2008), perhaps through disinhibition (Forstmann et al., 2008). Complementary research suggests that slower decisions involve inhibition from the subthalamic nucleus (Frank, Scheres, & Sherman, 2007; Aron & Poldrack, 2006). In either case, neurons encode a decision variable that, in a distributed manner, represents the evidence in favor of the decision. While these models are not strictly distributed control models (because the thresholding is assumed to be separate from the accumulation), they have characteristics of it. A major goal of the stopping literature is to identify the key brain site that regulates stopping. The distributed control approach cautions that such a site may need not exist; instead of a site, there might be a neural mechanism at work, one that is not distinct from the sites of neurons that form the perception-action stream.

### Conflict detection and resolution: honeybees

Humans performing a cognitively demanding task may realize the task is harder than expected and devote more effort to it (Botvinick et al., 1999; Shenhav, Botvinick, & Cohen, 2013). For example, in the Stroop task, a subject is asked to either read a word or name the color of ink used to display it (Stroop, 1935). Reading the word is easy, but naming the ink color, especially when the word itself is a different color, requires more cognitive control. Another form of conflict is decisional conflict, which is caused by ambivalence between two equally desired options (Cai & Padoa-Schioppa, 2012; Hayden, Heilbronner, & Pearson, 2011; Strait, Blanchard, & Hayden, 2014; Amiez, Joseph, and Procyk, 2006). Modular models of conflict detection and resolution generally involve a discrete conflict detector and resolver, which are often located in the dorsal anterior cingulate cortex (dACC, Botvinick et al., 1999, Shenhav, Botvinick, & Cohen, 2013; Botvinick et al., 2001). We hasten to note that such models, especially with regard to dACC, are contentious: the signal may not be conflict per se, but in either case, it may regulate control, which is our interest here (Kolling et al., 2016; Shenhav et al., 2016; Ebitz & Platt, 2015).

In springtime, thriving honeybee beehives reproduce. Roughly a third of the hive’s members remain at the hive site and the others leave to form a swarm that gathers in one location and, in a few days, chooses a new hive site from a radius of several kilometers (Seeley, 2010; Seeley & Burhman, 1999; Camazine et al., 1999). Like our ants above, scouts evaluate promising nearby sites and then return and signal their quality with special dances (Figure 7). Dances indicating higher quality sites induce other bees to investigate the same site. When scouts detect a quorum of bees at a site (typically around 20), they then return and provide a different signal, one that initiates a selection of the hive site by the swarm (Seeley, 2010; Seeley& Buhrman, 1999).

**Figure 7.**
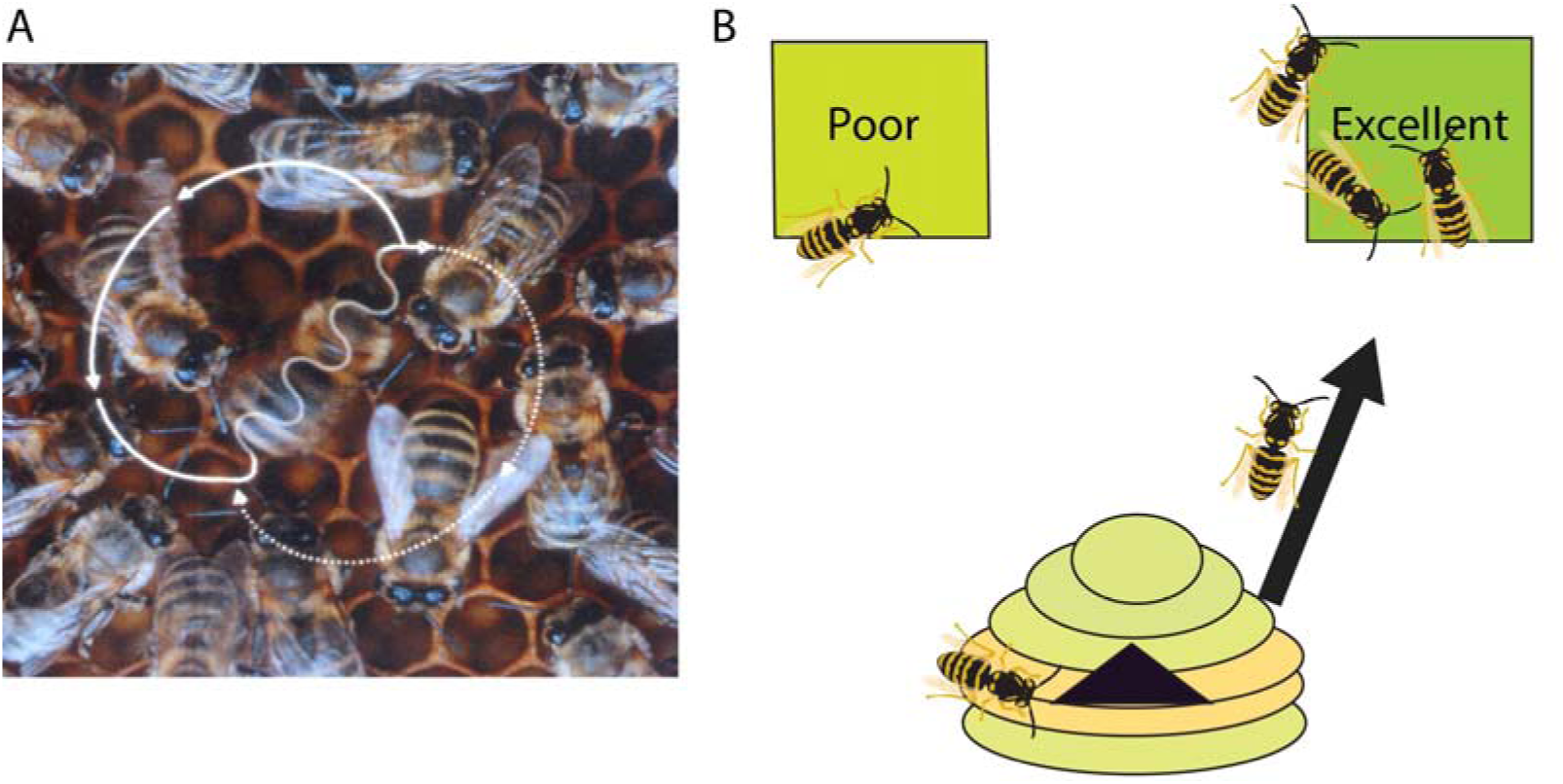
A) Image of honeybee waggle dance communication in a hive. Reproduced with permission from “Dances as a window into insect perception” (Chittka, 2004). B) Illustration of binary choice between hive sites. Through quorum sensing by scouts at potential nest cites and waggle dance communication with the swarm, new hive locations are efficiently chosen.

If there is one obvious best site, the decision will proceed quickly. But if there are two or more sites of approximately equal quality, the decision will proceed more slowly as the bees take the time to choose the best one. The swarm therefore is sensitive to decisional conflict: it monitors its own level of decisional ambivalence as the decision proceeds. Note that this is conflict signal a swarm, not individual variable; after all, no bee knows about more than one site, so no bee is conflicted. By not halting the search process, the swarm effectively recruits more processing resources (i.e. more bee-search time) when conflict is high. As in mental effort, deliberation is not free; swarms are vulnerable to weather and predators so there is an opportunity cost to delaying the construction of the hive (Lindauer, 1957).

Notably, the detection and resolution of conflict are emergent phenomena. No single bee that is sensitive to the conflict level – we know the rules the bees follow and none of them deal with conflict. Nor is there a conflict signal represented in the bee’s waggle dance or at any other point in the system. No bee has a specialized role before the swarm starts swarming. Still, the swarm as a whole is quite sensitive to decisional conflict and able to deal with it efficiently. It’s also worth noting that an aggregate measure of hive activity, say, the number of active scouts or number of active dances during the decision, will show clear and strong aggregate conflict signals. This finding is intriguing because conflict signals are seldom observed in the activity of single neurons, and yet are robustly observed in the brain’s hemodynamic activity (see below).

## Part IV. Evidence for distributed executive control in the brain

We turn now to the neuroscience of control. As noted above, there is a broad consensus that executive control is modular, not distributed (Botvinick & Cohen, 2014). We believe that one reason for relative unpopularity of distributed control systems by scholars is that they are unfamiliar and unintuitive. Indeed, distributed control is notoriously difficult for us to intuit. Terms like the “ghost in the machine,” “the invisible hand of the marketplace,” “asking the hive mind” are reminders that our own minds naturally impute discrete and coherent agency even when dealing with mindless and ghostless distributed systems. Still, many distributed control systems are intuitive and can become more so with familiarity.

### Neuroscience methods make modularity easier to find

Another factor disfavoring distributed control models is that the major methods for studying executive control, lesion, neuroimaging, and single unit recording, all arguably have some bias towards finding evidence of modularity.

Neuroimaging, like lesion studies, measures aggregate function of a given brain area or voxel, and thus cannot determine properties of the individual agents of the nervous system, neurons. This is true for multi-voxel pattern analysis as well as for ROI-type analyses. By aggregating signals across voxels, neuroimagers lose information about activity of individual neurons. The aggregate signal in turn misses information about the specific types of local, horizontal, and narrow-bandwidth signals that are crucial for distributed systems. But it is very good at detecting even weak signals at a broad range, meaning it can readily measure emergent properties of neural populations. The limitations of the lesion method are illustrated in a study by Plaut (1995). In this work, he shows how even the double dissociation, the gold standard of lesion studies, is susceptible to false positives supporting a modular view given certain reasonable assumptions about distributed network implementations of cognitive functions.

Single unit physiology studies are just as limited, although in the opposite way. Neurons may function much like agents, but the power of distributed systems comes in the specific local interactions of small numbers of agents. Physiology can measure the activity of only one neuron at a time; even multi-cellular methods have difficulty capturing interacting neurons. Moreover, most studies focus on a single brain region with the cost of inability to measure function at the level of the interregional network.

Historically, Karl Lashley had difficulty in finding the locus of memory function by lesion techniques (Lashley, 1929). This may have been because lesions to distributed systems do not selectively impair discrete functions, but instead have complex and unpredictable effects (Farah, 2004). Lashley found that degradation of behavioral performance depended on the amount of the brain regions removed independent of the precise location: they characteristically led to graceful degradation, which he interpreted as the product of mass action (Lashley, 1929). But when there is even a moderate amount of specialization in the system, they can lead to moderate but measurable effects. The interpretation of these effects, however, will be influenced by the experimenter’s theoretical framework.

### A case study: the dACC

To look at these general issues in detail, we will take the dorsal anterior cingulate (dACC, often just ACC) as a case study. The dACC is part of the cingulum, a band of cortex that wraps around the corpus callosum in the sagittal plane. The dACC receives a broad and diverse set of inputs that includes both limbic and cognitive regions, as well as dopamine signals, and projects to motor, premotor, and executive regions. These factors make it a natural site for serving as a monitor and controller. Indeed, a great deal of evidence links it to these two functions, among others. This evidence includes physiology (Heilbronner & Hayden, 2016), neuroimaging (Ridderinkoff et al., 2004; Shenhav, Botvinick, & Chohen, 2013; Kolling et al., 2012, Hare et al., 2011; Behrens et al., 2007; Hayden & Heilbronner, 2014), and lesion studies (Rudebeck et al., 2006; Kennerley et al., 2006; Picton et al., 2007; Turken & Swick, 1999). Most prominently its responses are activated by contexts that promote control (Rushworth et al., 2011; Shenhav, Botvinick, & Cohen, 2013). These include conflict (Botvinick et al., 19999; Ebitz & Platt, 2015; Sheth et al., 2012; but see Kolling et al., 2016 and Ebitz & Hayden, 2016), surprising and unexpected outcomes (Matsumoto et al., 2007; Hayden et al., 2011; Ito et al., 2003), rewards (Seo & Lee, 2007; Hayden, Pearson, & Platt, 2009; Kennerley et al., 2009); progression through a task (Ma et al., 2014; Shidara & Richmond, 2002; Hayden, Pearson, & Platt, 2011b), changes in environmental context and volatility (Behrens et al., 2007; Procyk, Tanaka, & Joeseph, 20000; Shima & Tanji, 1998), control of actions (Strait et al., 2016; Nakamura, Roesch, &Olson, 2005), and others not listed here. It is also directly activated by factors related to control, such as persistence (Blanchard, Strait, & Hayden, 2015; Chudasama et al., 2013; Parvizi et al., 2013; Hillman & Bilkey, 2012; Hillman & Bilkey, 2010).

These findings generally support a modular view of cognition, one in which dACC takes on the specialist role of monitor and controller. However, a broader review suggests that dACC is neither uniquely involved in monitoring and control, not is its function primarily these two roles. Indeed, the very long list of functions above should raise suspicion for a devotee of the modular viewpoint. Yes, these variables can all be placed under the rubric of monitoring and control, but at some point the definition becomes so elastic that it contains almost all of cognition. Second, are all these functions found only in the dACC? Unlikely. Most of these functions are shared with many other brain regions (Cisek & Kalaska, 2010). For example, recent work points to the important of the orbital surface in classically anterior cingulate functions like conflict monitoring and resolution (Mansouri, 2014), and regulating the explore-exploit tradeoff (Blanchard et al., 2015).

Studies that compare dACC activity with other brain regions often find that differences are more qualitative than quantitative (Hokosawa et al., 2013; Kennerley et al., 2009; Azab & Hayden, 2016). Indeed, control is associated with many other prefrontal structures, including OFC, dlPFC, vmPFC, and vlPFC (e.g. Schoenbaum et al., 2009; Wilson et al., 2014; Bechara, 2005; Buckley et al., 2009). Nor are these functions limited to the PFC; control signals are observed in the parietal cortex, the posterior cingulate cortex, the thalamus, and the striatum (e.g. Hayden, Smith, & Platt, 2010).

More broadly, summaries of dACC function tend to emphasize its potentially specialized role as a hub, linking visceral, cognitive, and motor systems (Bush, Luu, &Posner, 2000; Morecraft & VanHoesen, 1997; Rushworth et al., 2011; Paus, 2001; Heilbronner and Hayden, 2016). But is it really all that specialized? There is anatomical and functional evidence for it’s hub-nature, but it’s also true of other brain regions, including, for example, PCC (Heilbronner, Hayden, & Platt, 2011; Heilbronner & Platt, 2013) and insula. Indeed, rich interconnectivity is a feature of many brain systems (Wang & Kennedy, 2016; Heilbronner & Haber, 2014; Heilbronner et al., 2016).

Nor are the response properties observed in the dACC uniquely control-related. Many of them seem to fit naturally into the category of stimulus-response processing, rather than as a regulator of that processing. That is, if we think of the brain as a system that converts sensory inputs to motor outputs, we should expect in a modular brain to find no sensory and motor signals in dACC, and instead find pure control-selective signals (Cisek, 2012). Instead, dACC is prominently responsive to both sensory stimuli and to actions. One convenient parameter to look at is spatial representation; this is a prominent property of the physical world but should, in theory, not be part of the recondite world of control. And yet dACC encodes the locations of stimuli under consideration and the specific details of actions (Hayden & Platt, 2010; Isomura et al., 2003; Luk & Wallis, 2009; Stoll et al., 2016?; Strait et al., 2016; Shima & Tanji, 1998).

Together these pieces of evidence argue that the differences between the dACC and adjacent structures are not as strong as is conventionally believed. They suggest instead a broad continuity of function between dACC and its neighbors and afferents. The broad functions, especially in the control domain, that it serves, are more distributed than modular. Moreover, the units of dACC – its neurons – appear to play a role in input-output processing as well as in generation of control signals. That is, from the perspective of a scientist accustomed to thinking about bee swarms and ant colonies, they look much like individual bugs: sensitive to multiple task parameters and capable of generating their own control signals, which influence their neighbors, and have the capability of participating in a larger cascade and, under the right circumstances, having effects at the aggregate level.

### Maybe executive control could be distributed in the brain?

A priori, it is not unreasonable to think so. A basic description of the brain sounds like an ideal candidate for a distributed control system. Neurons are agents that can only communicate with a very small number of neighbors relative to the whole population. Like bacteria, they use a variety of diffusible chemicals to communicate. Each neuron can monitor an extremely limited portion of the world and can broadcast its signals to a very narrow part of the world as well. Each neuron has limited but powerful and non-linear computational properties.

Moreover, each cell is autonomous, but they work together, non-competitively, in the service of a much larger goal (overcoming competition is a major barrier for many distributed systems, Sumpter, 2006). Individual neurons possess the ability to regulate the activity of other neurons (or output structures) through changes in firing rate. This activity can serve as both a processing and a regulatory role. The properties of the whole system (the brain) are rich and flexible, much more so than any of its constituents (Hofstadter, 1985, Ch. 26). The brain makes use of both positive and negative feedback, and shows slow changes over time.

Strong circumstantial evidence for the distributed view comes from lesion studies (Farah, 2004; Wilson et al., 2010). Damage has surprisingly weak and graded effects; graceful degradation is a well-known property of distributed systems (McClelland et al., 1987). Of the major “clean” effects associated with lesions (prosopagnosia, hemianopia, scotoma, and so on), few would be considered executive control effects. Instead, impairments in executive control can come from lesions in many different areas, and associated effects are generally graded, and only grow serious when the lesions become quite large(Farah, 2004; Lashley, 1929; Wilson et al., 2010).

Although there is some evidence for control-specific lesions (Shallice, 1982; Levine et al., 1998; Duncan et al., 1996), it may be difficult to pin these data clearly to control functions. Instead, it may be that more difficult processing is impaired but simpler processing is spared. Consider, for example, an ant colony with a large proportion of members lesioned. That colony would have no trouble choosing a hive site if the decision was easy, but would have a great deal of trouble with a more difficult decision. We should not then conclude that the task-difficulty module is broken.

Indeed, the brain was the original inspiration for connectionist and PDP networks. The linkage between brain organization and other distributed control systems has been pointed out by many others before (Seeley, 2010; Couzin, 09; Passino et al., 2007; Mitchell, 2009). Given these facts, it is striking that the distributed view has not continued to serve as the null hypothesis for modular theories as a viable alternative view.

### Methods that can push for a distributed processing view

However, recent technological advances have made the distributed processing more attractive for researchers. With the adoption of newer analysis techniques, a host of traditional imaging methodologies are beginning to highlight the interconnectivity and coordination of many brain regions during a variety of tasks (Sporns and Betzel, 2016). For example functional connectivity analysis is a growing trend in fMRI imaging studies (Sporns and Betzel, 2016; Craddock, Tungaraza, and Milham, 2015). In contrast to traditional ROI analysis, functional connectivity analysis focuses on the interaction pattern between the brain regions as the determinant of brain function rather than the activity of the single brain regions (Craddock, Tungaraza, and Milham, 2015; Sporns, Tononi, and Kotter, 2005). Likewise, an increasing emphasis on large-scale brain networks has lead to a revision of cognitive functions extending across modular boundaries and sparked efforts to define functional regions based on “connectional fingerprints” (Misic & Sporns, 2016; Passingham, Stephan, & Kotter, 2002). These trends have lead to the new field of network analysis and connectomics that emphasizes the interconnections of different brain regions across structure and function. A recurrent theme in many studies utilizing network analysis is the distributed processing nature inherent to many tasks across brain regions over a singular key region (Wang et al., 2015; Bressler & Menon, 2010).

### The modular vs. distributed debate in stopping and working memory

For purposes of comparison, it is helpful to consider two aspects of executive control that have long been thought to be modular, but have more recently been challenged by a more distributed alternative view.

Influential work by Aron and others highlights the important and seemingly modular role of the right inferior frontal gyrus (rIFG) and anterior insula (aIns) in motor response inhibition, a form of executive control related to stopping (Rubia et al., 2001; Aron et al., 2003; Aron, Robbins, & Poldrack, 2004; Aron, Robbins, & Poldrack, 2014). However a recent series of studies challenges this view and proposes an alternative account that is more aligned with a distributed interpretation (Hampshire & Sharp, 2015; see also Munakata, 2011). Specifically, Hampshire and Sharp propose that stopping is the result of local processing by individual units that engage in lateral inhibition and potentiation, in a manner originally proposed for control of attention in the ventral stream (Desimone & Duncan, 1995; Chelazzi et al., 1998). In other words, they propose a simple set of local rules that neuron/agents can follow and produce effective stopping behavior. This view implements classic stopping models and is consistent with relevant unit physiology – that is, with measures of the responses of the putative agents (Band et al., 2003; Boucher et al., 2007; Schall, Stuphorn, & Brown, 2002). In contrast to Aron and colleagues, they propose that the rIFG/aIns is part of a larger multiple demand cortex that flexibly handles many executive functions, including stopping (Duncan, 2001; Cole & Schneider, 2007; Erika-Florence, Leech, & Hampshire, 2014). Ultimately, they suggest that stopping may not be a valid psychological construct, but rather a term used to describe intuitively similar behaviors.

Another example comes from the domain of working memory. Classic neurophysiological works by Niki, Fuster and then Goldman-Rakic supported the idea that the DLPFC serves as the site of working memory storage (Kubota & Niki, 1971; Funahashi, Bruce, & Goldman-Rakic, 1989; Alexander & Fuster, 1971; reviewed in Riley & Constantinidis, 2016). The key evidence for this idea was the fact that single neurons in that region showed systematic changes associated with the contents of working memory. This is a modular view: it proposes that specific rostral regions serve as sites of storage for working memory, while posterior regions implement perception and association. A recent body of work challenges this view and argues for a more distributed alternative (reviewed in Postle, 2006; Pasternak & Greenlea, 2005; Postle, 2016).

The alternative view proposes that neurons in frontal regions regulate storage (Lebedev et al., 2004; Postle, 2005), but that caudal regions responsible for perception are reactivated during working memory, and that their reactivation serves to store the information on-line (Harrison & Tong, 2009). This view thus sees perceptual neurons as flexible agents with multiple cognitive roles, including both basic processing and executive control roles. Indeed, further work suggests that modulations in these neurons may alter their responsiveness, thus serving as a form of proactive control that also implements memory-guided decisions (i.e. a matched filter, Machens, Romo, & Brody, 2005; Miller & Wang, 2006; David et al., 2008; Jun & Romo, 2010; Mirabella et al., 2007; Hayden & Gallant, 2013; Ogawa & Komatsu, 2004).

Working memory is interesting to use because of its centrality in the history of modular theories (i.e. most theories) of executive control (Baddeley Hitch, 1974; Baddeley, 1996). Especially, the concept of the central executive, which supports the short-term memory in demanding tasks, has been thought to play a diverse control functions. However, subsequent studies discredited the general function of the central executive and rather fractionated its functions to number of the different operations (Logie, 2016). Thus, as a psychological construct, the concept of the central executive in working memory might no longer be regarded as the modular, centralized function and rather as the functions of the distributed nature.

## CONCLUSIONS

We do not mean to imply that no current work could be classified as distributed. Quite the opposite is true. Many models have distributed aspects (e.g. Botvinick et al., 2001; OReily, Herd, & Pauli, 2010; Behrman & Plaut, 2013; Botvinick & Plaut, 2004; Munakata et al., 2010; Botvinick & Plaut, 2006; McClelland et al., 2010; Lenartowicz et al., 2010). Instead, our major goals are to highlight the key distinguishing features of distributed and modular systems.

### Advantages to a distributed control system

From the perspective of adaptiveness, there are several advantages of a distributed control system with simple agents (Brooks & Flynn, 1989). First, because it is self-organized, there is no need to build a special centralized organization system that will link up control elements with their corresponding processors. A modular system requires the equivalent of a telephone switchboard; a distributed one does not. Second, that self-organization gets around the specter of infinite regress (Cooper, 2010). For example, if we have a special centralized organization system, we need another system to build and maintain it, and to monitor its functioning, and so on, ad infinitum. Self-organizing systems are easier developmentally – there is no need to pre-specify their organization genetically or any other way. They are also more robust to damage and can more readily adapt and be amenable to plasticity, such as occurs with learning. They are generally more flexible for novel situations. Finally, and most important, distributed control is a good way to get complex and adaptive behavior from systems consisting of elements that are less complex (Sumpter, 2006). From a theoretical perspective, distributed system makes sense. Many brain functions are distributed, including perception and object recognition, storage of episodic memories, motor planning and execution, and, arguably, economic decision-making (Strait, Sleezer, & Hayden, 2015; Cisek, 2012; Cisek & Kalaska, 2010).

### How to study distributed executive control systems

Distributed control systems may be more difficult to study than modular ones with conventional methods. In many studies (including, we hasten to admit, many of our own), we pick out some psychological process of interest. We then ask whether brain activity in some neuron or voxel within a given brain region correlates with a measure of that variable. If we get a positive result, the simplest step is to infer that that variable is reified in the brain. The distributed perspective cautions against this strategy; such correlations may be real, but may only correlate with emergent properties of the system. And if the underlying processes are dissimilar, we will draw false conclusions. In other words, we are always in danger of reifying higher level processes at the lower level.

Instead, the best strategy for dealing with this possibility is a top-down research program. We should come up with specific hypotheses about how distributed control systems might work, and then estimate its expected neural signatures (e.g. Hampshire & Sharp, 2015). The next step is to identify the results expected from alternative distributed or even modular implementations, and perform the critical test of comparing alternative views. This approach is agnostic about method; it can be applied to unit physiology, neuroimaging, or even reaction times (Louie, Kaw, & Glimcher, 2013; Chau et al., 2014). And it’s worth reiterating that the two modular and the distributed views are not mutually incompatible. In reality, they may exist on a spectrum. And executive control may be heterogeneous; some aspects may be modular while others may be distributed.

The relevant hypotheses will come, as always, from close consideration of the data; especially from attempts to interpret data that conflict with preconceptions. But also, they can come from the animal kingdom, as we have discussed in this review. Brains are complex distributed systems, and they face many of the same constraints as others. It should not be surprising that they have a great deal in common with ant colonies, bee swarms, and herds of migrating baboons (Couzin, 2009; Sumpter, 206; Passino et al., 2007; Seeley, 2010; Hofstadter, 1980; Hofstadter, 1985, Ch. 25, 26).

## Acknowledgements

This work is supported by a CAREER award from NSF (BCS1253576) and a R01 from NIH (DA038615) to BYH. R.A. is supported by Research Fellowships for Young Scientists from the Japan Society for the Promotion of Science (JSPS). We thank Tom Seeley for patient explanations and Amanda Oglesby-Sherrouse for introducing us to the wonderful world of bioluminescent bacteria.

